# Kinesin-1 holoenzyme assembly coordinates cargo-adaptor recognition with heavy-chain autoinhibition

**DOI:** 10.64898/2026.08.20.746125

**Authors:** Jingling Niu, Mengmeng Zhang, Liping He, Xiaojie Zhu, Mingming Liu, Jiasheng Chen, Wenli Jiang, Chao Wang

## Abstract

Kinesin-1 is a major microtubule-based molecular motor that transports diverse cellular cargoes, yet how assembly of its subunits is coupled to cargo recognition and motor regulation remains poorly understood. Mammalian kinesin-1 functions as a heterotetrameric holoenzyme composed of kinesin heavy chains (KIF5s) and kinesin light chains (KLCs), but the structural principles linking holoenzyme assembly to motor regulation remain unclear. Here, using KIF5C as a model system, we define the molecular mechanisms underlying kinesin-1 holoenzyme assembly and cargo-adaptor regulation. We identify a conserved coiled-coil interface between KIF5C and KLC1 that mediates their 2:2 assembly, with quantitative mutagenesis revealing key hydrophobic determinants of complex formation. We further demonstrate that the mitochondrial adaptor TRAK2 directly engages the KIF5C CC4 cargo-binding platform through a defined 2:2 interaction required for mitochondrial recruitment of KIF5C. Although KLC1 and TRAK2 bind distinct regions of KIF5C, KLC1 modulates TRAK2 association through steric and conformational effects, revealing how holoenzyme composition influences cargo-adaptor accessibility. Mechanistically, we identify a previously unrecognized intramolecular interaction between the KIF5C CC1 and CC4 domains that forms a stalk-mediated autoinhibitory latch. TRAK2 and KLC1 release this inhibitory interaction through distinct mechanisms. Together, our study establishes a molecular framework in which kinesin-1 holoenzyme assembly regulates cargo-adaptor recognition and autoinhibitory remodeling, providing mechanistic insight into how molecular motors coordinate cargo engagement with activation.

## Introduction

Kinesin-1 is a major microtubule-based molecular motor that drives the long-range anterograde transport of diverse cellular cargoes, including mitochondria, endoplasmic reticulum (ER), Golgi-derived vesicles, and synaptic vesicle precursors, thereby supporting cellular homeostasis and neuronal function^1–5^. Unlike simple enzymatic machines, molecular motors must precisely coordinate motor activity with cargo engagement to ensure efficient and regulated intracellular transport. Mammalian kinesin-1 functions as a heterotetrameric holoenzyme composed of a homodimer of kinesin heavy chains (KIF5s) and two kinesin light chains (KLCs)^6–8^. Although extensive studies have elucidated the mechanochemical properties of KIF5 motor domains^9–11^ and the cargo-binding functions of KLCs^12–16^, how kinesin-1 holoenzyme assembly is organized at the molecular level and how this assembly influences cargo recognition and motor regulation remain poorly understood.

The kinesin-1 holoenzyme integrates distinct structural and regulatory modules contributed by KIF5 heavy chains and KLCs. The N-terminal motor domain of KIF5s couples ATP hydrolysis to conformational changes that drive directional movement along microtubules^17,18^. Following the motor domain, the elongated stalk region contains a series of coiled-coil domains that mediate heavy-chain dimerization and provide platforms for regulatory proteins and cargo adaptors. The neck coil CC0 contributes to heavy-chain homodimerization^19^. The central elbow region between CC2 and CC3 enables conformational flexibility required for compact folding^20–24^, whereas the C-terminal CC regions (CC3 and CC4) provide regulatory platforms for assembly with KLCs and engagement of cargo adaptors^25,26^. The disordered C-terminal tail contains a conserved IAK motif that participates in kinesin-1 autoinhibition by engaging the motor domains^27–30^. KLCs serve dual roles as structural components of the kinesin-1 holoenzyme and as cargo-recruiting adaptors. Through their N-terminal coiled-coil regions, KLCs associate with KIF5 heavy chains^25^, whereas their C-terminal six tetratricopeptide repeat (TPR) domains recognize diverse cargo adaptor proteins through short linear motifs, including W-acidic and related sequences^12,13,31–33^. In addition, an intervening linker region containing a conserved LFP motif can engage the TPR domains intramolecularly to regulate KLC conformation and cargo-binding activity^34^. Recent structural studies further suggest that KLC association contributes to stabilization of the compact kinesin-1 architecture and modulates the conformational state of the KIF5 motor complex^21,24,35^. However, the molecular basis governing KIF5-KLC assembly and the functional consequences of this interaction for kinesin-1 regulation remain elusive.

Cargo adaptor engagement provides a critical mechanism by which kinesin-1 activity is coupled to specific intracellular transport pathways. Cargo adaptors can interact with either KLCs or KIF5 heavy chains, and these interactions contribute to recruitment and regulation of kinesin-1 during cargo transport^36–39^. While cargo recognition mediated by KLC TPR domains has been extensively characterized, the molecular principles governing direct cargo-adaptor recognition by KIF5 heavy chains remain incompletely understood. Structural studies of Drosophila kinesin-1 revealed that the Kif5 CC4 region directly interacts with atypical Tropomyosin-1/C (aTm1) through an antiparallel heterotrimeric coiled-coil assembly, providing one of the few structural examples of heavy-chain-mediated cargo recognition^40,41^. Whether similar mechanisms operate in mammalian kinesin-1 and how KIF5-mediated cargo recognition is coordinated with holoenzyme assembly remain unclear.

The trafficking kinesin-binding proteins TRAK1 and TRAK2 represent physiologically important cargo adaptors that connect kinesin-1 and dynein–dynactin complexes with Miro-associated mitochondria to regulate bidirectional mitochondrial transport^42–46^. TRAK proteins contain N-terminal coiled-coil regions responsible for motor recruitment and activation^47,48^, as well as C-terminal regions that interact with Miro proteins on the mitochondrial surface^49,50^. Although TRAK proteins are known to associate with kinesin-1 and promote mitochondrial transport, the molecular basis underlying TRAK-KIF5 recognition and how this interaction is coordinated with KLC-mediated holoenzyme assembly remain unknown. Moreover, although the conserved IAK motif contributes to kinesin-1 autoinhibition, disruption of this canonical interaction does not fully abolish the inhibited state, suggesting the existence of additional regulatory elements within the kinesin-1 architecture^21,51^. Understanding how holoenzyme assembly, cargo adaptor binding, and heavy-chain autoinhibition are coordinated is therefore essential for elucidating the molecular principles governing kinesin-1 activation.

Here, we used KIF5C as a model system to define how kinesin-1 holoenzyme assembly regulates cargo-adaptor recognition and autoinhibition. Through biochemical, structural modeling, and cellular approaches, we characterized the molecular basis of KIF5C association with KLC1 and identified the principles governing heavy chain-light chain assembly. We further defined the interaction between KIF5C and the mitochondrial cargo adaptor TRAK2 and revealed how KLC1 modulates TRAK2 engagement with KIF5C. In addition, we identified a previously unrecognized intramolecular interaction between the KIF5C CC1 and CC4 domains that contributes to stalk-mediated autoinhibition. Notably, both TRAK2 binding and KLC1 association relieve this inhibitory interaction through distinct mechanisms, linking holoenzyme assembly and cargo recognition to autoinhibitory regulation. Our findings establish a mechanistic framework in which kinesin-1 holoenzyme assembly, cargo-adaptor recognition, and heavy-chain autoinhibition are coupled to regulate the transition between inactive and cargo-engaged states.

## Results

### KIF5C and KLC1 assemble through a defined 2:2 coiled-coil interface

Mammalian kinesin-1 functions as a heterotetrameric holoenzyme composed of KIF5 heavy chains and kinesin light chains (KLCs), yet the molecular basis governing heavy chain-light chain assembly remains poorly defined. Previous structural studies have suggested that KLCs contribute to the compact architecture and regulatory state of kinesin-1^21,24,35^, but the molecular interface and organization principles underlying KIF5-KLC assembly remain unclear (Fig. 1A). To define the molecular basis of kinesin-1 holoenzyme assembly, we systematically characterized the interaction between KIF5C and KLC1 using diverse biochemical approaches. Co-immunoprecipitation (Co-IP) assays in HEK-293T cells revealed that KLC1 selectively associates with the KIF5C CC3 region, whereas other regions of KIF5C showed no detectable interaction (Fig. 1B). Reciprocal Co-IP analyses further demonstrated that KIF5C binding is mediated by the N-terminal region of KLC1, while the C-terminal tetratricopeptide repeat (TPR) domain, which mediates cargo adaptor recognition, is dispensable for heavy chain association (Fig. 1C). Purified protein assays using fast protein liquid chromatography (FPLC) and isothermal titration calorimetry (ITC) confirmed that the KIF5C CC3 domain directly engages the KLC1 N-terminal region with submicromolar affinity (Supplementary Fig. 1). These results define a specific assembly interface between KIF5C and KLC1 and distinguish the heavy chain-binding function of KLC1 from its cargo-recruiting activity.

**Figure 1.**
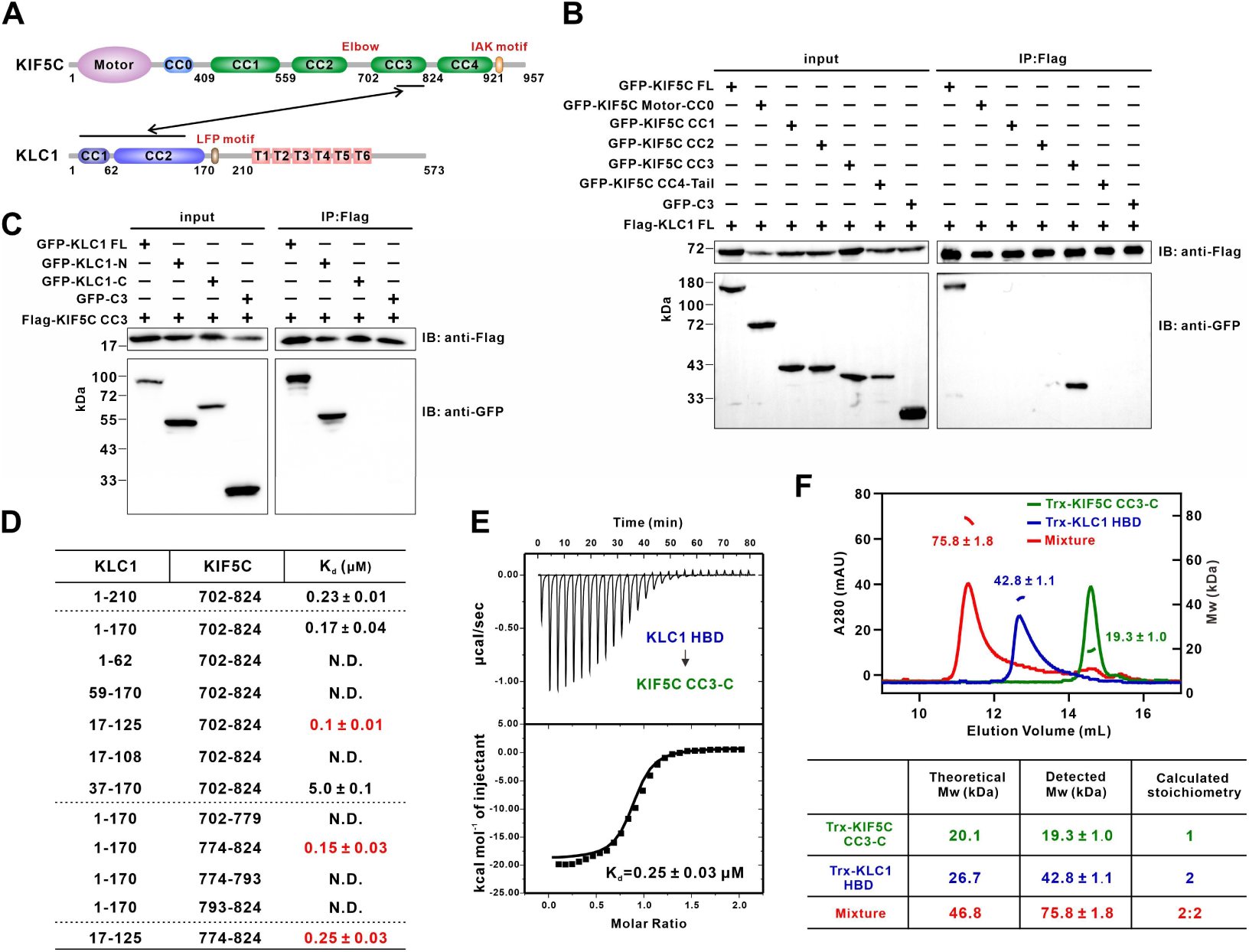
Molecular basis of KIF5C-KLC1 holoenzyme assembly. **(A)** Schematic representation of the domain organizations of KIF5C and KLC1. The identified intermolecular assembly interface between KIF5C CC3 and the KLC1 N-terminal heavy chain-binding domain (HBD) is indicated. CC, coiled-coil domain; T1-T6, tetratricopeptide repeat domains 1-6. **(B)** Co-immunoprecipitation (Co-IP) analysis of the interaction between KLC1 and KIF5C truncation variants in HEK-293T cells. Flag-KLC1 full-length (FL) was immunoprecipitated, and associated GFP-tagged KIF5C fragments were detected by immunoblotting. KLC1 selectively associated with the KIF5C CC3 region. **(C)** Reciprocal Co-IP assays defining the KLC1 region responsible for KIF5C binding. Flag-KIF5C CC3 was co-expressed with GFP-tagged KLC1 full-length (FL), N-terminal region (KLC1-N, residues 1-210), or C-terminal TPR-containing region (KLC1-C, residues 211-573). The KIF5C-binding activity was mediated by the KLC1 N-terminal region, whereas the TPR domain was dispensable for heavy chain association. **(D)** Mapping of the minimal KIF5C-KLC1 assembly interface. ITC assays using KLC1 N-terminal truncation variants and KIF5C CC3 fragments identified KLC1 residues 17-125 as the minimal heavy chain-binding domain (HBD) and KIF5C residues 774-824 within CC3 as the minimal KLC1-binding region (CC3-C). N.D., no binding detected. **(E)** ITC measurement of the direct interaction between purified KIF5C CC3-C and KLC1 HBD. The binding affinity was determined by fitting the data to a one-site binding model. The reported K_d_ value represents the fitting result obtained using Origin 7.0. **(F)** Fast protein liquid chromatography coupled with multi-angle light scattering (FPLC-MALS) analysis defining the oligomeric state and stoichiometry of the KIF5C CC3-C/KLC1 HBD complex. The experimentally determined molecular weight is consistent with a 2:2 stoichiometry.

To further resolve the molecular determinants of this interaction, we performed systematic truncation mapping of both binding partners. The intact tandem coiled-coil region within the KLC1 N-terminus was required for KIF5C association, as individual KLC1 CC1 or CC2 domains failed to support detectable binding (Fig. 1D, rows 2-4). Further refinement identified KLC1 residues 17-125 as the minimal heavy chain-binding domain (HBD) (Fig. 1D, rows 5-7). Conversely, analysis of KIF5C truncations demonstrated that the C-terminal portion of CC3 (CC3-C, residues 774-824) is necessary and sufficient for KLC1 binding (Fig. 1D, rows 8-11). ITC assays confirmed that the isolated KIF5C CC3-C and KLC1 HBD directly interact with a K_d_ of approximately 0.25 μM (Fig. 1E). Moreover, FPLC coupled with multi-angle light scattering (FPLC-MALS) demonstrated that the KIF5C CC3-C/KLC1 HBD complex adopts a 2:2 stoichiometry (Fig. 1F), consistent with the organization of the native kinesin-1 holoenzyme. Together, these results define a coiled-coil assembly module between KIF5C and KLC1 and provide a molecular framework for kinesin-1 holoenzyme formation.

### Conserved hydrophobic layers stabilize the KIF5C-KLC1 assembly

Having defined the minimal KIF5C-KLC1 assembly interface, we next sought to identify the structural principles that stabilize this interaction. Despite extensive crystallization screening using multiple KIF5C and KLC1 truncation constructs, including co-purified complexes and engineered fusion proteins, we were unable to obtain crystals suitable for structure determination. We therefore used AlphaFold3 to model the KIF5C CC3-C and KLC1 HBD complex. The predicted structure revealed a symmetrical six-helix bundle in which two KLC1 molecules dimerize through their CC2 regions, while two KIF5C molecules pack against the KLC1 dimeric core to form a compact coiled-coil assembly (Figs. 2A and 2B).

**Figure 2.**
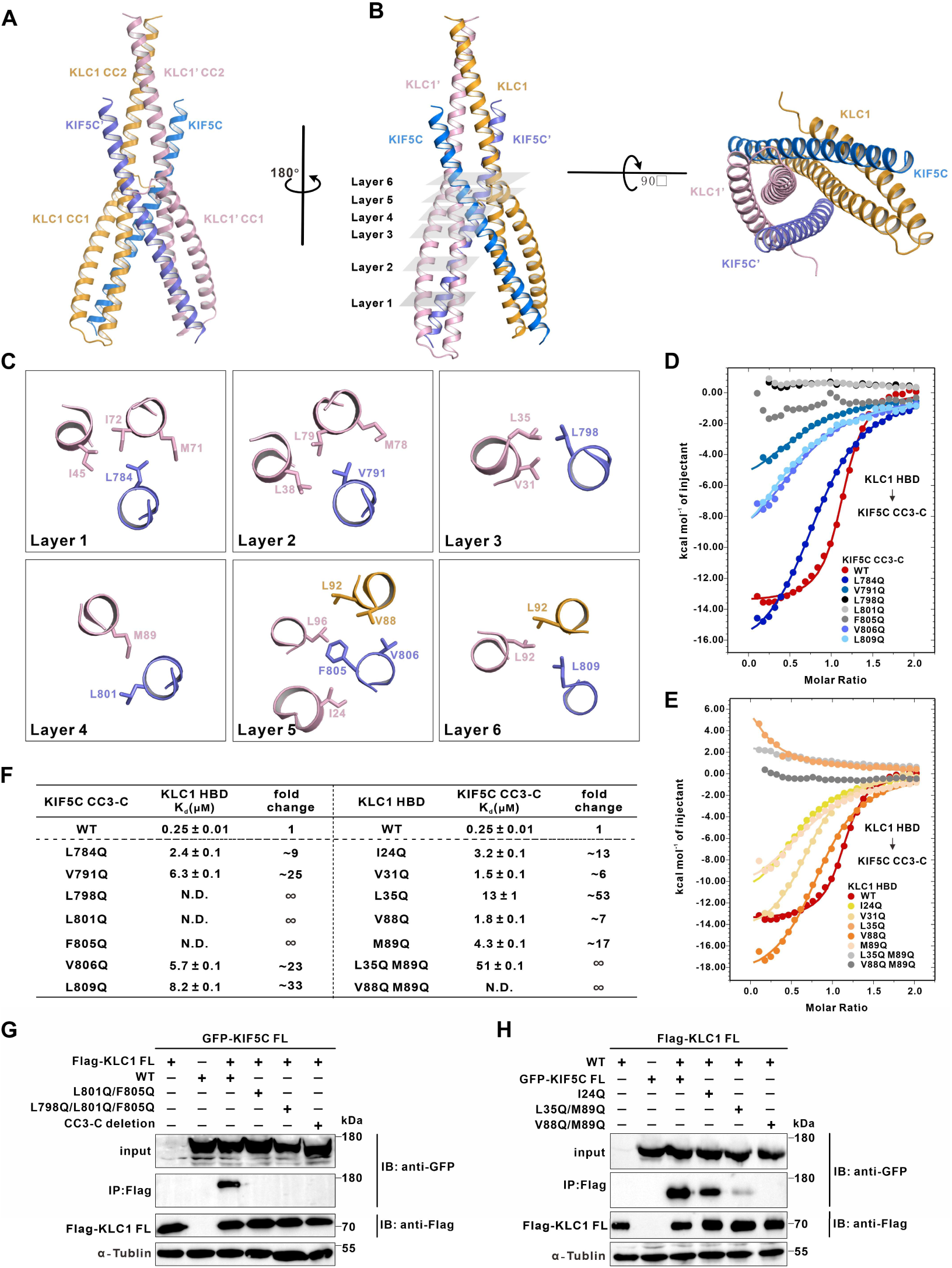
Conserved hydrophobic layers stabilize KIF5C-KLC1 holoenzyme assembly. **(A)** Ribbon representation of the AlphaFold3-predicted KIF5C CC3-C and KLC1 HBD complex. The two KIF5C molecules are shown in marine blue and light blue, and the two KLC1 molecules are shown in orange and deep salmon. **(B)** Overall architecture of the predicted KIF5C-KLC1 assembly interface. Six stacked hydrophobic layers formed by interacting side chains are indicated along the coiled-coil bundle. **(C)** Close-up views of the six hydrophobic layers at the KIF5C–KLC1 interface. Key side chains involved in hydrophobic packing are shown as sticks and labeled. **(D)** ITC-derived binding curves measuring the interaction between KLC1 HBD and wild-type KIF5C CC3-C or the indicated KIF5C mutants (L784Q, V791Q, L798Q, L801Q, F805Q, V806Q, and L809Q). **(E)** ITC-derived binding curves measuring the interaction between KIF5C CC3-C and wild-type KLC1 HBD or the indicated KLC1 mutants (I24Q, V31Q, L35Q, V88Q, M89Q, L35Q/M89Q, and V88Q/M89Q). **(F)** Summary of the ITC-derived K_d_ values for the KIF5C and KLC1 mutants shown in (D) and (E), illustrating the mutational effects on KIF5C-KLC1 binding. **(G)** Co-IP analysis of full-length KIF5C mutants with KLC1 in HEK-293T cells. The L801Q/F805Q double mutation, the L798Q/L801Q/F805Q triple mutation, and deletion of the CC3-C region disrupted KIF5C-KLC1 interaction. α-Tubulin served as a loading control. **(H)** Reciprocal Co-IP analysis of full-length KLC1 interface mutants with KIF5C in HEK-293T cells. Mutations targeting the hydrophobic assembly interface impaired KLC1 association with KIF5C, with combined mutations (L35Q/M89Q and V88Q/M89Q) producing near-complete or complete loss of interaction, respectively. α-Tubulin served as a loading control.

Analysis of the predicted interface suggested that the KIF5C-KLC1 assembly is stabilized predominantly by hydrophobic interactions, with only limited contribution from electrostatic contacts (Supplementary Fig. 2). Specifically, nonpolar side chains from neighboring helices interlock to generate six stacked hydrophobic layers across the interface (Figs. 2B and 2C). To test this model, we introduced glutamine substitutions into the corresponding hydrophobic residues of KIF5C and quantified binding to KLC1 by ITC. Mutations across the interface weakened the interaction to varying extents, with KIF5C L798Q and L801Q in layers 3 and 4, as well as F805Q in layer 5, showing the strongest effects (Figs. 2D and 2F). Structural analysis indicated that KIF5C L798 and L801 pack against KLC1 V31/L35 and M89, respectively, whereas KIF5C F805 and V806 are buried within an extensive hydrophobic cleft formed by residues from both KLC1 subunits (Fig. 2C), highlighting layer 5 as a major stabilizing element. We next examined the reciprocal contribution of the KLC1 interface. Mutations targeting hydrophobic residues within layers 3-5 consistently impaired KIF5C binding. Among these residues, KLC1 L35Q in layer 3 caused a greater than 50-fold reduction in binding affinity, whereas individual substitutions of V88Q or M89Q within layers 4 and 5 weakened the interaction. Notably, combined disruption of these hydrophobic contacts, including the L35Q/M89Q and V88Q/M89Q double mutations, nearly abolished or completely abolished KIF5C binding, respectively (Figs. 2E and 2F), demonstrating that multiple hydrophobic layers cooperatively stabilize the KIF5C–KLC1 interface.

To validate the predicted interface in the context of full-length proteins, we introduced layer-specific mutations into both KIF5C and KLC1 and examined their interaction by reciprocal Co-IP assays. Consistent with the biochemical measurements, individual mutations in KIF5C, including L798Q, L801Q, and F805Q, weakened but did not eliminate KLC1 association (Supplementary Fig. 3), whereas the L801Q/F805Q double mutation and the L798Q/L801Q/F805Q triple mutation completely disrupted KIF5C-KLC1 interaction (Fig. 2G). Conversely, mutations introduced into the corresponding hydrophobic residues of full-length KLC1 similarly impaired its association with KIF5C, with combined interface mutations producing the strongest defects (Fig. 2H). Together, these reciprocal mutational analyses validate the predicted assembly model and demonstrate that kinesin-1 holoenzyme formation is driven by a cooperative hydrophobic coiled-coil interface involving multiple layers of intermolecular packing.

### The mitochondrial adaptor TRAK2 directly engages the KIF5C CC4 cargo-binding platform

With the molecular basis of KIF5C-KLC1 holoenzyme assembly established, we next sought to determine how kinesin-1 recognizes physiological cargo adaptors through the KIF5 heavy chain. We focused on TRAK2, a mitochondrial adaptor protein that links kinesin-1 to Miro-associated mitochondria and regulates mitochondrial transport. TRAK2 contains N-terminal coiled-coil regions responsible for motor association and a C-terminal region involved in mitochondrial recruitment (Fig. 3A).

**Figure 3.**
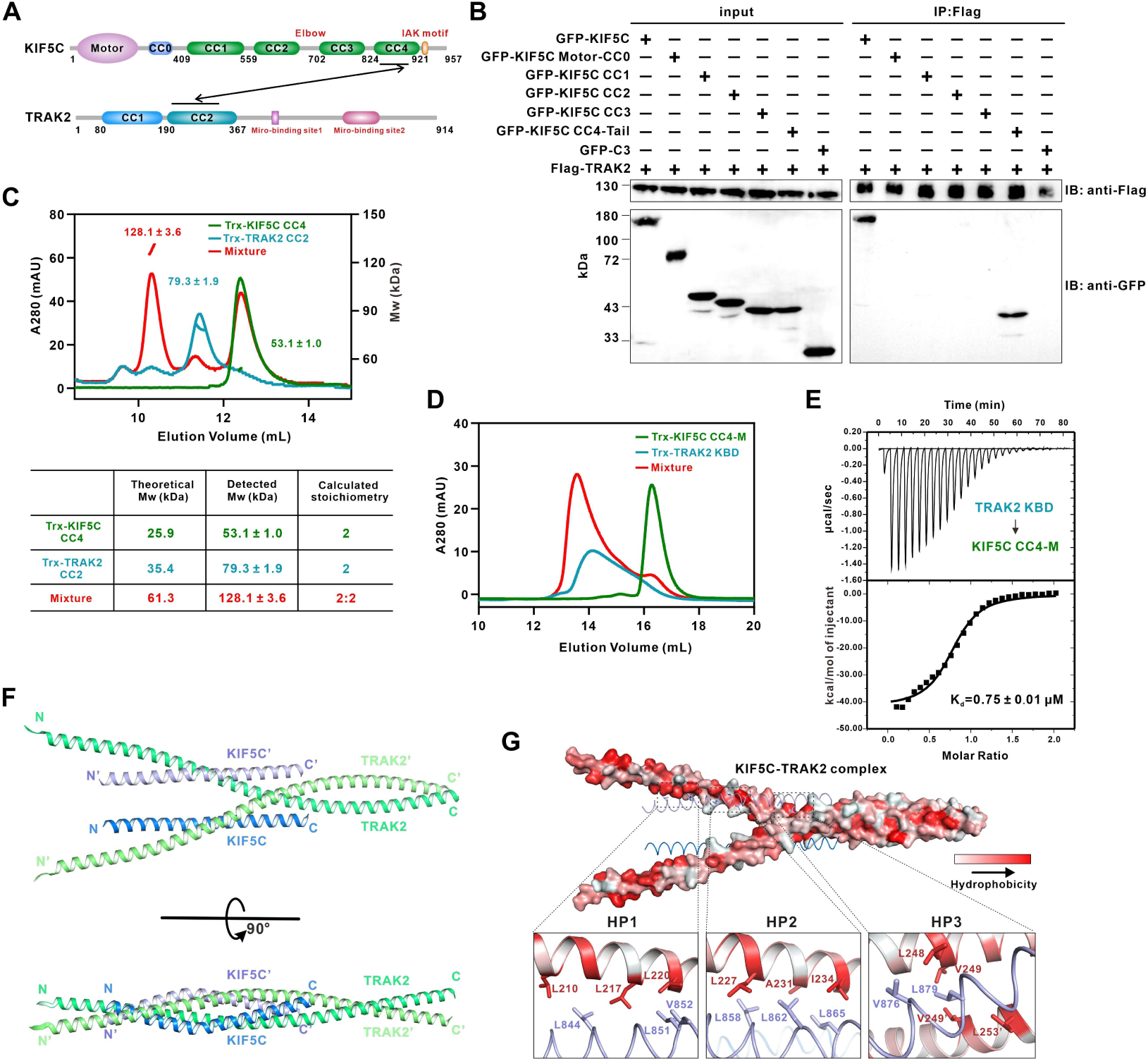
TRAK2 directly engages the KIF5C CC4 cargo-binding platform. **(A)** Schematic representation of the domain organizations of KIF5C and TRAK2. The minimal interaction regions identified between KIF5C and TRAK2 are indicated. **(B)** Co-IP analysis of the interaction between TRAK2 and KIF5C truncation variants in HEK-293T cells. TRAK2 selectively associated with the KIF5C CC4-Tail region. **(C)** FPLC-MALS analysis defining the oligomeric states and stoichiometry of KIF5C CC4, TRAK2 CC2, and the KIF5C-TRAK2 complex. The calculated and experimentally measured molecular weights are summarized in the table. **(D, E)** FPLC (D) and ITC (E) analyses demonstrating the direct interaction between KIF5C CC4-M (residues 838-890) and TRAK2 KBD (residues 190-301). ITC analysis was used to determine the binding affinity of this interaction. **(F)** Ribbon representation of the AlphaFold3-predicted TRAK2 KBD and KIF5C CC4-M complex. KIF5C molecules are shown in marine blue and light blue, whereas TRAK2 molecules are shown in dark green and light green. **(G)** Hydrophobic surface representation of the TRAK2-KIF5C interface. Three hydrophobic patches (HP1-3) are highlighted, and close-up views show the residues involved in intermolecular packing.

Co-IP screening in HEK-293T cells revealed that TRAK2 selectively associates with the CC4-Tail region of KIF5C, whereas other KIF5C regions showed no detectable interaction (Fig. 3B). Given that CC4 represents a major cargo-binding platform within KIF5s, we purified KIF5C CC4 together with the two N-terminal coiled-coil domains of TRAK2. FPLC-MALS analysis demonstrated that TRAK2 CC2 directly associates with KIF5C CC4 with a 2:2 stoichiometry, consistent with the dimeric organization of both proteins (Fig. 3C), whereas TRAK2 CC1 exhibited no detectable interaction (Supplementary Fig. 4). Further biochemical mapping using FPLC and ITC identified TRAK2 residues 190-301 as the kinesin-1 binding domain (KBD) and KIF5C residues 838-890 within CC4 (CC4-M) as the minimal interaction regions, which directly interact with a K_d_ of approximately 0.75 μM (Figs. 3D and 3E).

To define the structural basis of this interaction, we generated an AlphaFold3 model of the TRAK2 KBD and KIF5C CC4-M complex. The predicted structure revealed a parallel four-helix bundle in which the TRAK2 dimer adopts an open-scissors-like conformation, allowing two KIF5C CC4 molecules to engage independently with the two arms of the adaptor dimer (Fig. 3F). Detailed analysis of the interface identified three hydrophobic patches (HP1-3) that define the TRAK2-KIF5C recognition interface. In HP1, KIF5C L844, L851, and V852 interact with TRAK2 L210, L217, and L220. HP2 is formed by interactions between KIF5C L858, L862, and L865 and TRAK2 L227, A231, and I234. At HP3, KIF5C V876 and L879 insert into a hydrophobic pocket formed by residues within the TRAK2 dimer interface (Fig. 3G). Together, these results define the molecular basis by which the mitochondrial adaptor TRAK2 directly engages the KIF5C CC4 cargo-binding platform.

### TRAK2-KIF5C interface mutations disrupt cargo adaptor-mediated mitochondrial recruitment of KIF5C

Having defined the structural basis of the TRAK2-KIF5C interaction, we next performed quantitative mutational analysis to validate the predicted interface and determine the functional importance of individual contact sites. Mutations targeting the HP1 interface consistently weakened TRAK2-KIF5C binding. In particular, the KIF5C L851Q/V852Q and TRAK2 L217Q/L220Q double mutations reduced binding affinity by more than 10-fold (Figs. 4A and 4D). The HP2 and HP3 interfaces contributed more substantially to complex stability. Within HP2, the KIF5C L858Q/L862Q double mutation completely abolished detectable binding, whereas the TRAK2 L227Q mutation severely impaired the interaction, resulting in a loss of measurable binding by ITC (Figs. 4B and 4D). Similarly, disruption of HP3 by KIF5C L879Q or TRAK2 L248Q/V249Q mutations eliminated detectable binding (Figs. 4C and 4D). Together, these quantitative binding analyses validate the contribution of the predicted hydrophobic interface to TRAK2-KIF5C recognition.

**Figure 4.**
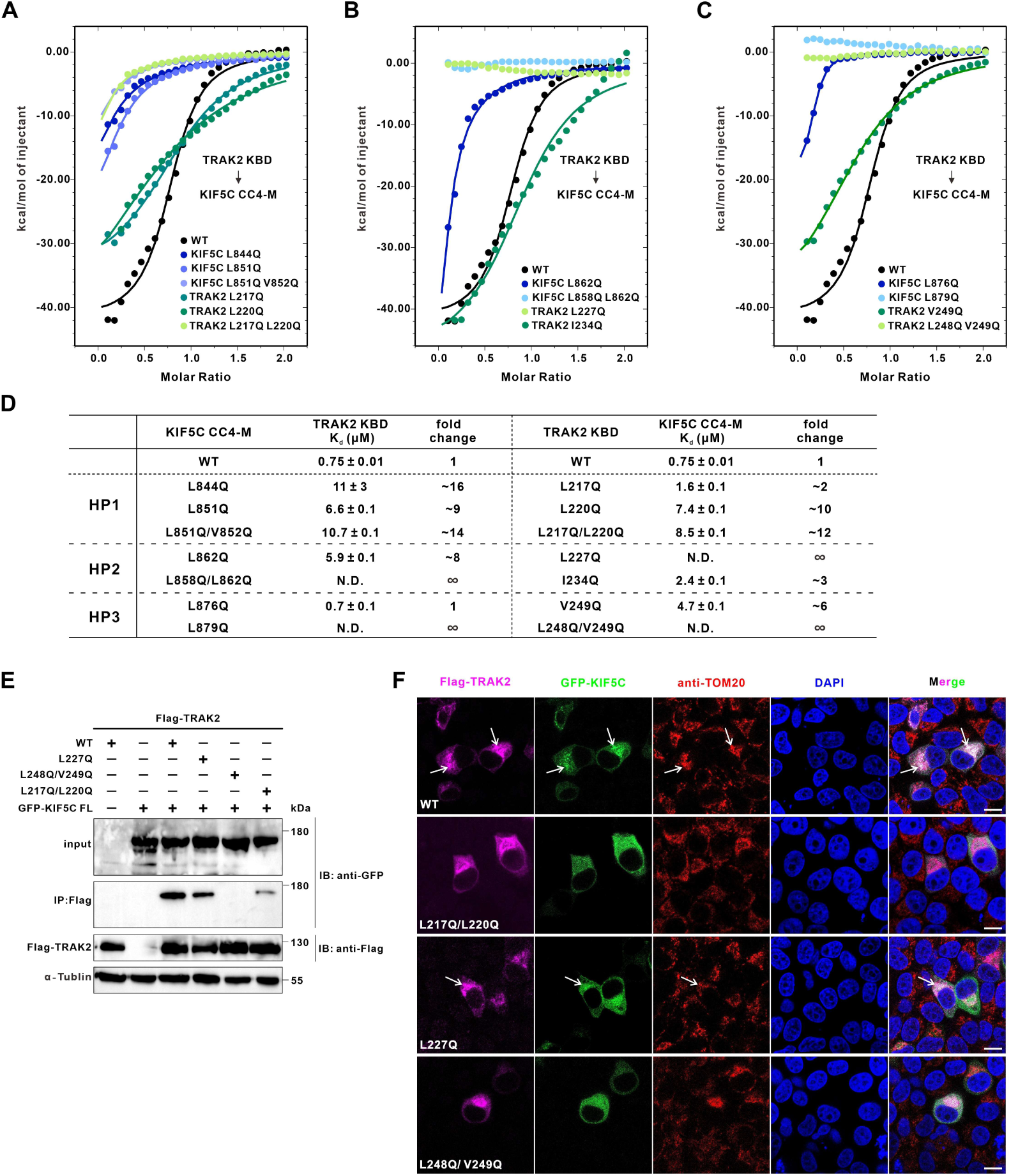
TRAK2–KIF5C interface mutations disrupt adaptor-mediated mitochondrial recruitment of KIF5C. **(A)** ITC analysis of the HP1 interface between KIF5C and TRAK2. Binding curves were obtained using wild-type or indicated KIF5C/TRAK2 mutants. **(B)** ITC analysis of the HP2 interface between KIF5C and TRAK2. Binding curves were obtained using wild-type or indicated KIF5C/TRAK2 mutants. **(C)** ITC analysis of the HP3 interface between KIF5C and TRAK2. Binding curves were obtained using wild-type or indicated KIF5C/TRAK2 mutants. **(D)** Summary of the ITC-derived K_d_ values from (A)-(C). The K_d_ values of various TRAK2 or KIF5C mutants are compared with their respective WT counterparts to highlight the mutational effects on the binding affinity. **(E)** Co-IP analysis of the interaction between full-length KIF5C and TRAK2 mutants in HEK-293T cells. TRAK2 L248Q/V249Q abolished KIF5C association, whereas L217Q/L220Q markedly reduced binding. TRAK2 L227Q retained partial association with full-length KIF5C. α-Tubulin served as a loading control. **(F)** Immunofluorescence analysis of KIF5C mitochondrial recruitment mediated by WT or mutant TRAK2 in HeLa cells. GFP-KIF5C (green) and Flag-TRAK2 (magenta) were visualized together with TOM20-positive mitochondria (red). Mutations disrupting TRAK2-KIF5C binding impaired mitochondrial enrichment of KIF5C without affecting TRAK2 mitochondrial localization. Nuclei were stained with DAPI. White arrows indicate mitochondrial-localized Flag-TRAK2. Scale bar, 10 μm.

We next examined whether these interface mutations affected the interaction and recruitment of full-length proteins in cells. Co-IP assays demonstrated that TRAK2 L248Q/V249Q completely abolished association with full-length KIF5C, whereas TRAK2 L217Q/L220Q markedly reduced KIF5C binding. In contrast, TRAK2 L227Q retained partial association with full-length KIF5C (Fig. 4E), suggesting that additional contacts within the intact proteins contribute to complex formation. TRAK2 recruits kinesin-1 to mitochondria to mediate mitochondrial transport. To determine whether disruption of the TRAK2–KIF5C interface affects mitochondrial recruitment, we examined the localization of GFP-KIF5C in HeLa cells co-expressing WT or mutant TRAK2. Wild-type TRAK2 strongly colocalized with GFP-KIF5C and promoted its enrichment on TOM20-positive mitochondria. Importantly, TRAK2 mutations that disrupted KIF5C binding did not affect TRAK2 mitochondrial localization itself. However, TRAK2 L217Q/L220Q and L248Q/V249Q mutants failed to recruit KIF5C to mitochondria, resulting in diffuse cytosolic distribution of GFP-KIF5C. Consistent with the residual full-length interaction observed by Co-IP, the TRAK2 L227Q mutant retained partial capacity to enrich KIF5C on mitochondria (Fig. 4F). Together, these findings validate the functional importance of the TRAK2-KIF5C interface for adaptor-mediated mitochondrial recruitment of kinesin-1 and reveal a direct heavy-chain-mediated cargo adaptor recognition mechanism underlying cargo engagement.

### KLC1 regulates TRAK2 engagement with KIF5C through structural occlusion and conformational remodeling

Previous studies have suggested that KLC modulates KIF5-mediated mitochondrial transport by regulating cargo adaptor engagement^52^. Consistent with this notion, fluorescence microscopy revealed that KLC1 expression reduced the colocalization between KIF5C and TRAK2 in mammalian cells (Fig. 5A). However, our biochemical analyses demonstrated that KLC1 and TRAK2 recognize distinct, non-overlapping regions of KIF5C (Figs. 1 and 3), indicating that KLC1-mediated inhibition of TRAK2 recruitment is unlikely to result from direct competition for the same binding site. We therefore sought to determine how KLC1 regulates TRAK2 association with the KIF5C heavy chain.

**Figure 5.**
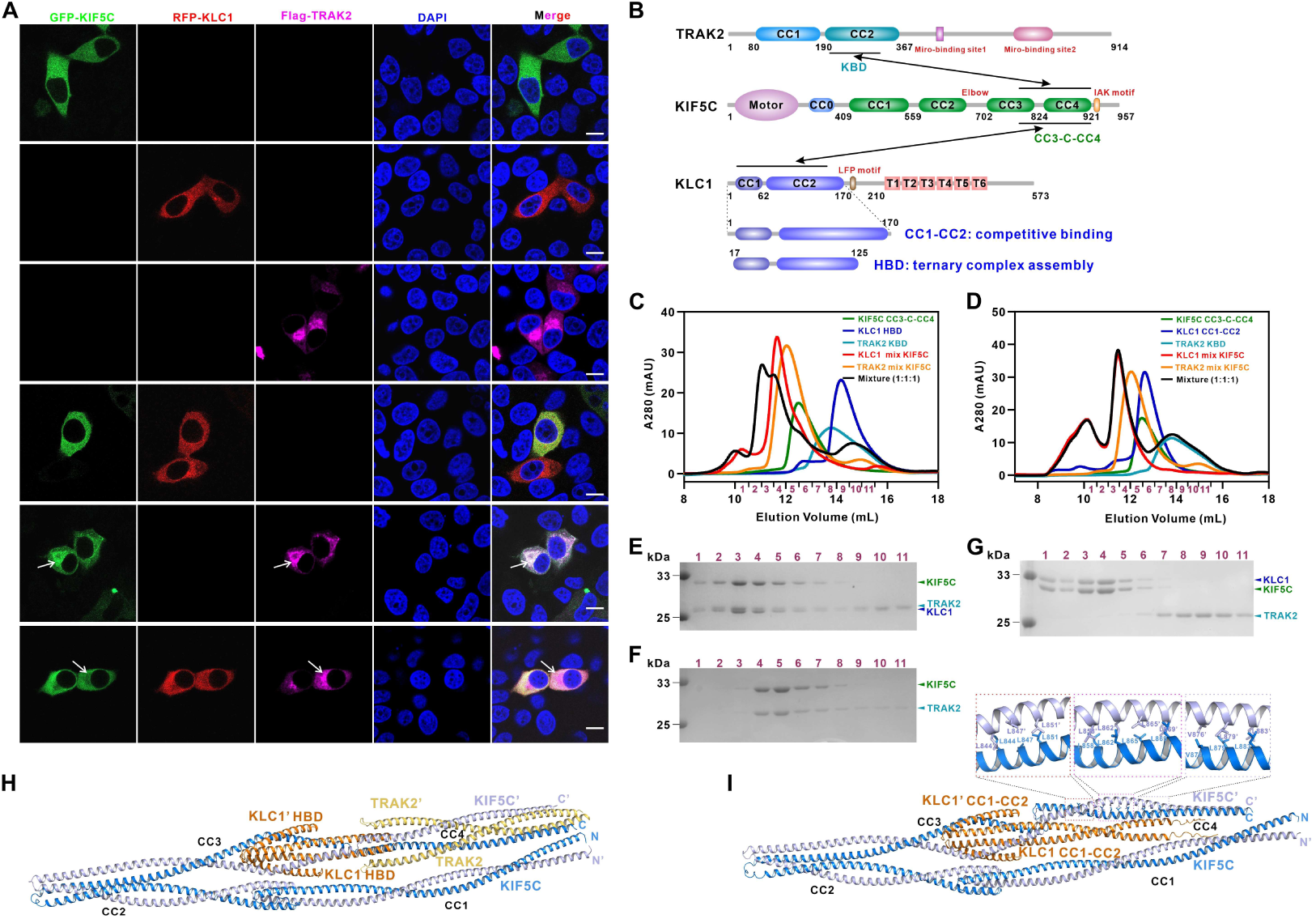
KLC1 regulates TRAK2 engagement with KIF5C through structural occlusion and conformational remodeling. **(A)** Representative immunofluorescence images showing intracellular localization and colocalization of KIF5C, KLC1, and TRAK2 in HeLa cells. Cells were transfected with GFP-KIF5C (green), RFP-KLC1 (red), and Flag-TRAK2 (magenta; detected by anti-Flag antibody), individually or in combination as indicated. White arrows indicate mitochondrial-localized Flag-TRAK2. Scale bar, 10 μm. **(B)** Domain organization of TRAK2, KIF5C, and KLC1, with interacting regions indicated. KLC1 truncation constructs used in this study are shown below. The KLC1 HBD fragment (residues 17-125) permits ternary complex formation with KIF5C and TRAK2, whereas the extended KLC1 CC1-CC2 fragment (residues 1-170) displaces TRAK2 from KIF5C. **(C)** FPLC analysis of complex formation among KIF5C, KLC1 HBD, and TRAK2. Binary mixtures of KIF5C with either KLC1 HBD or TRAK2 exhibited shifted elution profiles compared with individual proteins. The ternary mixture showed an additional shift, indicating formation of a stable KIF5C–KLC1 HBD–TRAK2 complex. Fractions analyzed by subsequent SDS-PAGE are indicated. **(D)** FPLC analysis of complex formation among KIF5C, KLC1 CC1-CC2, and TRAK2. The elution profile of the ternary mixture overlapped with that of the KIF5C-KLC1 CC1-CC2 binary complex, indicating that extended KLC1 CC1-CC2 competes with TRAK2 for KIF5C engagement. Fractions analyzed by subsequent SDS-PAGE are indicated. **(E, F)** SDS-PAGE analysis of FPLC fractions from panel (C). Coomassie staining of the ternary KIF5C-TRAK2-KLC1 HBD complex showing co-elution of all three components (E). KIF5C-TRAK2 binary complex as control (F). **(G)** SDS-PAGE analysis of FPLC fractions from panel (D). KLC1 CC1-CC2 and KIF5C co-elute in early fractions, whereas TRAK2 is excluded from the complex and elutes separately. **(H)** AlphaFold3-predicted structure of the ternary complex consisting of KIF5C CC1-CC4 (residues 409-921; marine blue and light blue), KLC1 HBD (residues 17-125; orange), and TRAK2 KBD (residues 190-301; yellow). The model illustrates that KLC1 HBD does not obstruct the TRAK2-binding platform on KIF5C CC4. **(I)** AlphaFold3-predicted structure of the KIF5C CC1-CC4-KLC1 CC1-CC2 complex. The extended C-terminal helix of KLC1 inserts between the CC1 and CC4 domains, occluding TRAK2 docking. Enlarged views highlight conformational rearrangement of CC4 and redistribution of hydrophobic residues involved in TRAK2 recognition toward CC4 self-association.

To dissect the molecular basis of this regulation, we generated a KIF5C fragment encompassing residues 774-921 (designated CC3-C-CC4), which contains both the CC4 domain and the major KLC1-binding region (Fig. 5B). This KIF5C fragment was sufficient to independently interact with either TRAK2 or KLC1, allowing us to examine their potential interplay. Surprisingly, when KIF5C CC3-C-CC4 was incubated simultaneously with TRAK2 and the KLC1 HBD fragment (residues 17-125), the three proteins assembled into a stable ternary complex (Figs. 5C, 5E, and 5F). This result indicates that the minimal KLC1-binding module does not interfere with TRAK2 recognition of the KIF5C heavy chain. In contrast, inclusion of an extended KLC1 construct containing residues 1-170 (designated CC1-CC2) completely displaced TRAK2 from the KIF5C complex (Figs. 5D, 5F, and 5G). Given that the extreme N-terminal region of KLC1 is intrinsically disordered, these findings suggest that the extended helical region within KLC1 CC1-CC2 provides the structural element required to restrict TRAK2 engagement. We therefore hypothesized that this extension may either physically obstruct the TRAK2 docking platform or induce conformational rearrangements within KIF5C that compromise TRAK2 recognition.

Structural modeling supported this hypothesis. In the predicted ternary complex, the shorter KLC1 HBD fragment binds to KIF5C without contacting the CC4 domain, leaving the CC4 platform accessible for TRAK2 engagement. Accordingly, the two arms of TRAK2 can simultaneously interact with the two CC4 regions of the KIF5C homodimer, supporting our structural model that TRAK2 binding remodels the local CC4 self-association interface (Fig. 5H). By contrast, in the KIF5C-KLC1 CC1-CC2 complex, the extended C-terminal helix of KLC1 occupies the space between the CC1 and CC4 domains, creating steric hindrance that blocks access of one TRAK2 docking arm to the CC4 platform (Fig. 5I). Moreover, KLC1 CC1-CC2 binding induces a conformational rearrangement of CC4, redirecting key hydrophobic residues previously identified as critical determinants of TRAK2 recognition, including L844, L851, L858, L862, and L879, toward CC4 self-association (Fig. 5I). Together, these findings reveal that KLC1 regulates cargo adaptor accessibility through structural remodeling of the KIF5C heavy chain rather than direct competition for the TRAK2-binding interface, providing a molecular mechanism by which kinesin-1 assembly is coordinated with cargo adaptor recruitment.

### KIF5C CC1 engages CC4 to establish an intramolecular autoinhibitory latch

The ability of KLC1 to regulate cargo adaptor engagement raised a fundamental question: how does cargo adaptor binding overcome the intrinsically inactive conformation of kinesin-1? We therefore sought to identify the structural elements within KIF5C that maintain its autoinhibited state. Through a systematic Co-IP analysis of interactions among KIF5C functional domains, we found that the compact intramolecular architecture of KIF5C is maintained by two distinct interaction interfaces: the canonical Motor-CC0/CC4-Tail interaction and a previously unrecognized CC1-CC4 interaction (Figs. 6A-6C). In contrast, no detectable interactions were observed between the CC2 or CC3 domains and other structural modules (Supplementary Fig. 5), suggesting that these regions primarily function as flexible spacers within the kinesin stalk rather than as direct mediators of autoinhibitory contacts.

**Figure 6.**
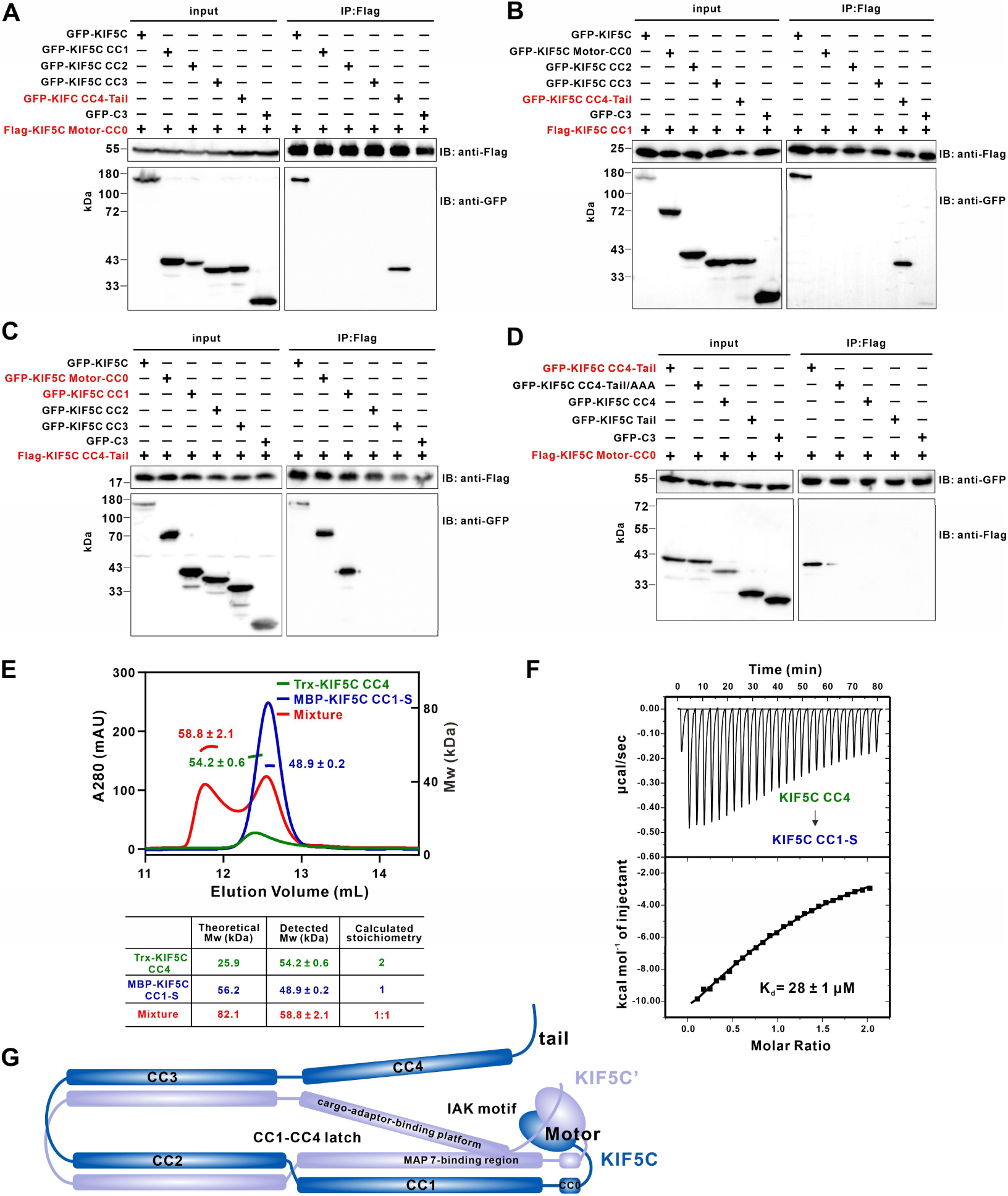
Identification of a CC1-CC4 intramolecular latch that maintains KIF5C autoinhibition. **(A-C)** Co-IP analysis of the intramolecular interaction network among KIF5C functional domains in HeLa cells. Flag-tagged KIF5C Motor-CC0 (A) or CC1 (B) was co-expressed with indicated GFP-tagged KIF5C fragments. Reciprocal Co-IP was performed using Flag-tagged KIF5C CC4-Tail (C). These analyses identify two intramolecular interaction interfaces within KIF5C: the canonical Motor-CC0/CC4-Tail interaction and the CC1-CC4 interaction. **(D)** Co-IP analysis of the canonical tail-mediated autoinhibitory interface. Mutation of the conserved IAK motif to AAA disrupts the Motor-CC0/CC4-Tail interaction. Motor-CC0 does not independently interact with isolated CC4 or Tail domains, suggesting that proper tail context is required for motor docking. **(E)** FPLC-MALS analysis of the oligomeric states and complex stoichiometry of KIF5C CC4 and CC1-S. CC4 forms a homodimer alone, whereas CC1-S binding disrupts CC4 self-association and generates a 1:1 CC1-S-CC4 complex. Molecular weights determined experimentally and theoretically are summarized. **(F)** ITC measurement of the direct interaction between KIF5C CC1-S and CC4. The CC1-S-CC4 interaction exhibits moderate affinity with a measured Kd of approximately 28 μM. **(G)** Schematic model illustrating how the CC1-CC4 intramolecular latch may cooperate with canonical tail–motor docking to stabilize the autoinhibited conformation of KIF5C.

To validate the canonical tail-mediated autoinhibitory interaction, we introduced mutations into the conserved IAK motif within the KIF5C tail region. Substitution of the IAK motif with AAA almost completely abolished the interaction between Motor-CC0 and CC4-Tail (Fig. 6D), confirming that this interface is mediated by the established tail-motor autoinhibitory mechanism. Interestingly, Motor-CC0 alone failed to interact with either isolated CC4 or the tail domain (Fig. 6D). We reasoned that truncation of the tail region surrounding the IAK motif may disrupt the structural context required for proper docking of the tail into the motor domain. We next investigated the newly identified CC1-CC4 interaction. Purification of KIF5C CC4 and a truncated CC1 fragment (residues 439-534, designated CC1-S) enabled direct biochemical characterization of this interface. FPLC-MALS analysis revealed that CC4 exists as a homodimer alone, whereas addition of CC1-S disrupted the CC4 homodimer and generated a 1:1 CC1-S-CC4 complex (Fig. 6E). Consistently, ITC measurements demonstrated a direct interaction between CC1-S and CC4 with a moderate affinity (K_d_ ≈ 28 μM) (Fig. 6F).

Structural considerations provide insight into how this interaction may contribute to kinesin-1 autoinhibition. The tail region containing the IAK motif immediately follows CC4, and a pair of motor domains asymmetrically engages the tail to establish the canonical inhibited conformation^27–30^. Within the elongated kinesin stalk, the CC1-CC4 interaction may constrain stalk flexibility and organize the spatial arrangement of the downstream tail, thereby facilitating efficient tail docking onto the motor domains (Fig. 6G). Moreover, because CC1 serves as a microtubule-associated protein 7 (MAP7)-binding region^53–55^, whereas CC4 functions as a cargo adaptor-binding platform, their intramolecular association may simultaneously restrict access to both regulatory surfaces, preventing premature engagement with microtubules or cargo molecules (Fig. 6G). Together, these findings identify a previously unrecognized CC1-CC4 intramolecular latch within KIF5C that cooperates with the canonical tail-motor interaction to stabilize the autoinhibited kinesin-1 conformation and provides a structural framework for adaptor-mediated activation.

### TRAK2 and KLC1 dismantle the CC1-CC4 autoinhibitory latch through distinct mechanisms

The identification of the CC1-CC4 interaction as an autoinhibitory latch raised the question of how this inhibitory contact is released during kinesin-1 activation. Because the CC4 domain serves as a shared binding platform for both CC1 and TRAK2, and because TRAK2 binds CC4 with substantially higher affinity than CC1 (Figs. 3E and 6F), we hypothesized that TRAK2 may directly relieve the CC1-CC4 latch through competitive binding. To test this idea, we mixed KIF5C CC1-S, KIF5C CC4, and TRAK2 KBD *in vitro* and analyzed the resulting complexes by FPLC and SDS-PAGE. TRAK2 efficiently displaced CC1-S from CC4, resulting in formation of the TRAK2-CC4 complex and release of free CC1-S (Fig. 7A-7D). These results indicate that TRAK2 directly dismantles the CC1-CC4 autoinhibitory latch by competitively engaging the shared CC4 binding surface.

**Figure 7.**
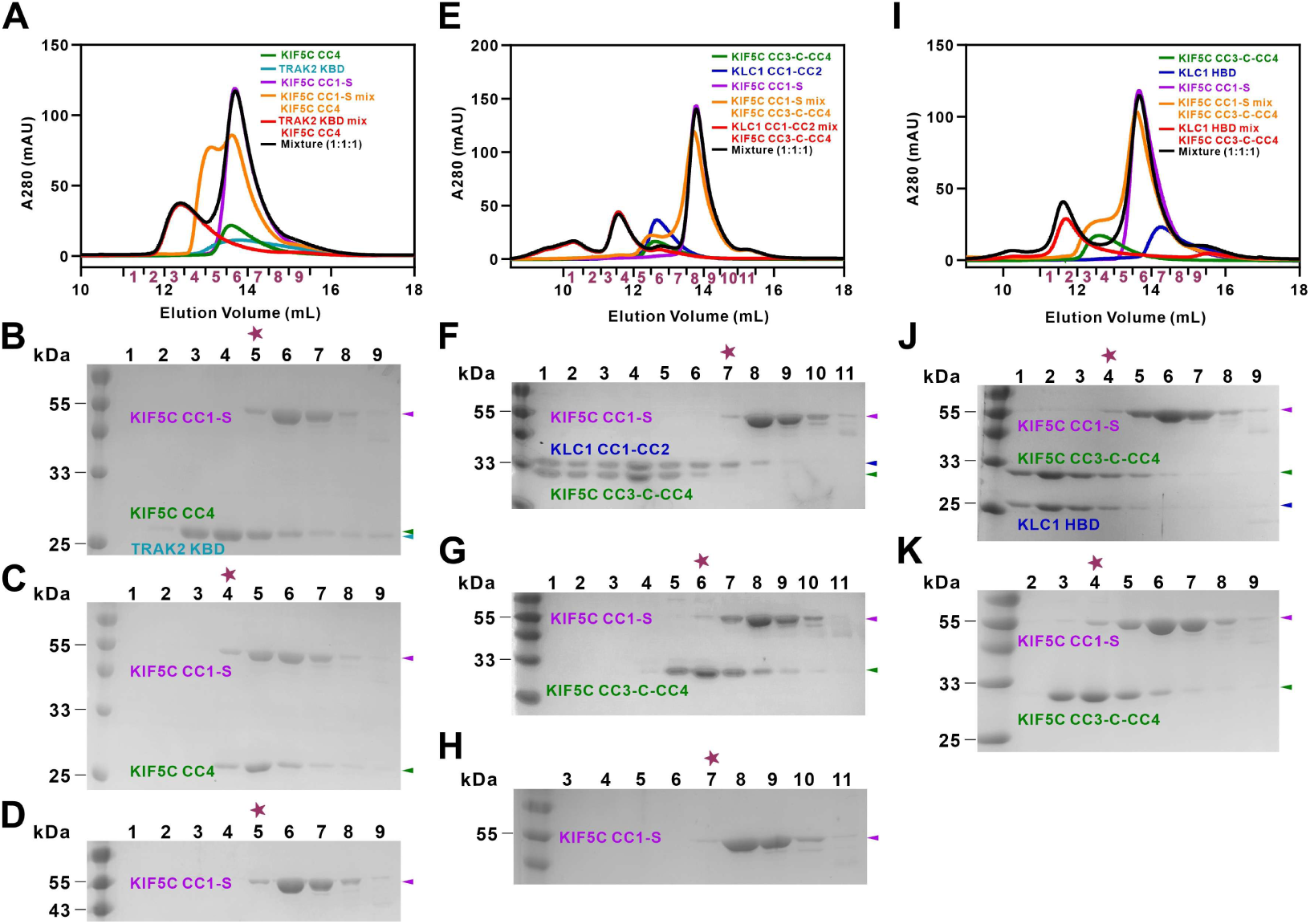
TRAK2 and KLC1 dismantle the CC1-CC4 autoinhibitory latch through distinct mechanisms. **(A)** FPLC analysis of the interaction among KIF5C CC1-S, KIF5C CC4, and TRAK2 KBD. Fractions analyzed by subsequent SDS-PAGE are indicated. **(B-D)** SDS-PAGE analysis of the fractions shown in panel (A). Coomassie-stained gels of the 1:1:1 mixture containing KIF5C CC1-S, KIF5C CC4, and TRAK2 KBD (B); KIF5C CC1-S and KIF5C CC4 binary mixture (C); KIF5C CC1-S alone (D). Asterisks indicate the fractions in which free KIF5C CC1-S begins to elute. **(E)** FPLC analysis of the interaction among KIF5C CC1-S, KIF5C CC3-C-CC4, and KLC1 CC1-CC2. Fractions analyzed by subsequent SDS-PAGE are indicated. **(F-H)** SDS-PAGE analysis of the fractions shown in panel (E). Coomassie-stained gels of the 1:1:1 mixture containing KIF5C CC1-S, KIF5C CC3-C-CC4, and KLC1 CC1-CC2 (F); KIF5C CC3-C-CC4 and KIF5C CC1-S binary mixture (G); KIF5C CC1-S alone (H). Asterisks indicate the fractions in which free KIF5C CC1-S begins to elute. **(I)** FPLC analysis of the interaction among KIF5C CC1-S, KIF5C CC3-C-CC4, and KLC1 HBD. Fractions analyzed by subsequent SDS-PAGE are indicated. **(J, K)** SDS-PAGE analysis of the FPLC fractions shown in panel (I). Coomassie-stained gels show the protein elution profiles of the 1:1:1 mixture containing KIF5C CC1-S, KIF5C CC3-C-CC4, and KLC1 HBD (J); KIF5C CC1-S and KIF5C CC3-C-CC4 binary complex control (K). The asterisks indicate fractions in which free KIF5C CC1-S begins to elute.

We next considered how this inhibitory interaction might be relieved in kinesin-1 heterotetramers, in which cargo adaptors such as SKIP are recruited through the KLC1 TPR domains rather than through direct binding to KIF5C CC4. This prompted us to ask whether KLC1 itself could relieve the CC1-CC4 latch. As suggested by the KIF5C-KLC1 CC1-CC2 structural model (Fig. 5I), the extended C-terminal region of KLC1 inserts between the CC1 and CC4 segments of KIF5C, spatially separating these two elements and thereby potentially disrupting their intramolecular association. Consistent with this model, biochemical reconstitution showed that KLC1 CC1-CC2 binding to KIF5C CC3-C-CC4 effectively blocked its interaction with CC1-S *in vitro* (Fig. 7E-7H). In contrast, the shorter KLC1 HBD fragment, which lacks the extended C-terminal helix and does not impose steric separation, had no detectable effect on the CC1-CC4 interaction (Fig. 7I-7K). These findings indicate that KLC1 relieves the CC1-CC4 latch not by direct competition for the CC4 surface, but by sterically and structurally disrupting the intramolecular fold that maintains KIF5C autoinhibition.

Together, our results define two mechanistically distinct routes for release of the CC1-CC4 autoinhibitory latch. In KIF5C homodimers, the mitochondrial cargo adaptor TRAK2 directly unlocks the latch through high-affinity competitive binding to CC4. In kinesin-1 heterotetramers, KLC1 acts as a structural activation switch that dismantles the same inhibitory interaction through steric separation and conformational remodeling. These dual regulatory modes provide a molecular framework linking cargo adaptor recognition and holoenzyme assembly to activation of the kinesin-1 motor.

## Discussion

Growing evidence has revealed that kinesin-1 regulation extends beyond the classical tail-to-motor docking model and involves coordinated interactions among the motor domain, stalk, tail, and kinesin light chains^21,24,35,51^. In this study, we establish a comprehensive mechanistic framework describing how kinesin-1 assembly, cargo adaptor recognition, and autoinhibitory control are structurally integrated within the KIF5C heavy chain. We first define the molecular architecture underlying KIF5C-KLC1 holoenzyme assembly and identify a conserved hydrophobic coiled-coil interface that stabilizes the heterotetrameric motor complex. Beyond intermolecular assembly, we uncover a previously unrecognized intramolecular CC1-CC4 interaction within the KIF5C stalk that cooperates with the canonical tail-mediated inhibition to maintain the compact inactive state of kinesin-1. Together, these findings reveal a multilayered regulatory architecture in which the KIF5C heavy chain functions not merely as a motor scaffold, but as an integrated regulatory hub coordinating motor assembly, cargo recognition, and activation.

Cargo recognition represents a central step in kinesin-1 regulation, yet the molecular mechanisms by which distinct adaptors engage the heavy chain remain incompletely understood. Here, using the mitochondrial adaptor TRAK2 as a model, we demonstrate that the KIF5C CC4 domain serves as a direct cargo-binding platform that recruits a TRAK2 homodimer through a defined helical interaction interface. The open-scissors-like architecture of the TRAK2 dimer enables two KIF5C CC4 molecules to engage independently, generating a parallel four-helix bundle that differs substantially from previously characterized kinesin-cargo adaptor assemblies, such as the antiparallel heterotrimeric coiled-coil formed between KIF5 and aTm1 during *oskar* mRNA transport^40^. These structural differences highlight the versatility of the KIF5 heavy chain in recognizing diverse cargo adaptors through distinct molecular strategies. Importantly, disruption of the TRAK2–KIF5C interface abolished mitochondrial recruitment of KIF5C in cells, demonstrating that this direct heavy-chain-mediated interaction is functionally coupled to organelle transport.

Our findings further reveal an unexpected regulatory role for KLC1 in controlling cargo adaptor accessibility. Although KLC1 and TRAK2 bind spatially distinct regions of KIF5C, KLC1 imposes a selective gatekeeping mechanism that limits TRAK2 engagement through structural remodeling of the heavy chain rather than direct competition for the same binding site. The extended C-terminal region of KLC1 CC1-CC2 introduces steric constraints that reshape the CC4 region and alter the accessibility of the TRAK2-binding platform. This mechanism expands the current view of KLC1, which has traditionally been considered primarily as a cargo adaptor-binding scaffold through its TPR domains^12–14,31–33^. Instead, our results demonstrate that KLC1 also functions as an intrinsic conformational regulator of the kinesin-1 heavy chain, providing a structural link between holoenzyme assembly and cargo adaptor recognition.

The identification of the CC1-CC4 intramolecular latch provides a mechanistic explanation for how cargo engagement can be coupled to kinesin-1 activation. This newly defined interaction, together with the canonical tail–motor association, establishes a multilayered autoinhibitory architecture that stabilizes the inactive conformation of KIF5C. We show that TRAK2 and KLC1 can release this inhibitory latch through distinct mechanisms: TRAK2 directly competes for the shared CC4 binding surface with higher affinity, whereas KLC1 disrupts the interaction through steric separation and conformational remodeling. These findings suggest that kinesin-1 activation is not controlled by a single molecular switch, but rather by coordinated transitions among multiple structural states governed by cargo adaptor engagement and holoenzyme composition. Future structural studies of full-length kinesin-1 complexes captured in distinct functional states, particularly during microtubule engagement and cargo transport, will be essential to visualize the long-range conformational transitions connecting stalk remodeling, tail release, and motor activation.

Together, our study establishes the KIF5C heavy chain as an integrated regulatory hub that couples holoenzyme assembly, cargo adaptor recognition, and autoinhibitory control. This mechanistic framework provides new insights into how kinesin-1 achieves precise regulation of cargo-dependent transport and highlights the importance of coordinated structural transitions in the functional diversification of molecular motors.

## Materials and Methods

### Constructs, protein expression and purification

The coding sequences of human KIF5C (UniProt: O60282), KLC1 (UniProt: Q07866), and TRAK2 (UniProt: O60296) were PCR-amplified from a human brain cDNA library. Truncated fragments and targeted mutants of KIF5C, KLC1, and TRAK2 were generated using standard PCR-based methods, and all sequences were verified by DNA sequencing. For recombinant protein expression, selected coding sequences were subcloned into modified pET32a-derived vectors. N-terminal thioredoxin-His_6_-tagged or MBP-His_6_-tagged fusion proteins were expressed in *Escherichia coli* BL21 (DE3) cells cultured in LB medium at 16 °C. Recombinant proteins were initially purified using a Ni-NTA agarose affinity chromatography, followed by size-exclusion chromatography using a Superdex 200 pg column (26/600, GE Healthcare) equilibrated with buffer containing 50 mM Tris-HCl (pH 7.8), 100 mM or 300 mM NaCl, 1 mM EDTA, and 1 mM DTT.

### Fast protein liquid chromatography (FPLC) assay

Analytical gel filtration chromatography was performed on an AKTA pure system (GE Healthcare). Protein samples (70μM) were loaded onto a Superdex 200 Increase column (GE Healthcare) equilibrated with a buffer containing 50 mM Tris-HCl (pH 7.8), 100 mM or 300 mM NaCl, 1 mM EDTA, and 1 mM DTT. Elution profiles were analyzed and plotted using GraphPad Prism 8 (GraphPad Software).

### FPLC coupled with multi-angle light scattering (FPLC-MALS) assay

FPLC-MALS analysis was performed using an AKTA pure system (GE Healthcare) coupled with a multi-angle light scattering detector (miniDAWN, Wyatt) and a differential refractive index detector (Optilab, Wyatt). Protein samples (70 μM for KIF5C, KLC1, and TRAK2) were filtered and loaded onto a Superdex 200 Increase column (GE Healthcare). Molecular weights were calculated using ASTRA 7 software (Wyatt).

### Isothermal titration calorimetry (ITC) assay

Isothermal titration calorimetry (ITC) measurements were carried out on a VP-ITC MicroCal calorimeter (Malvern) at 25 °C. All protein samples were prepared in a buffer containing 50 mM Tris-HCl (pH 7.8), 100 mM or 300 mM NaCl, 1 mM EDTA, and 1 mM DTT. Protein samples at 200µM (KLC1 or TRAK2) were loaded into the syringe, and KIF5C fragments at 20µM were loaded into the cell. Each injection consisted of a 10 μL aliquot with an interval of 180 s to allow the heat signal to return to baseline. ITC data were analyzed using Origin 7.0 software and fitted using a one-site binding model.

### Structure prediction

Three-dimensional structures of protein complexes were predicted using AlphaFold3. The KIF5C-KLC1 complex, TRAK2-KIF5C complex, and KIF5C-KLC1-TRAK2 ternary complex were modeled independently. Default parameters were used, and five structural models were generated for each complex. The highest-ranked model was selected for subsequent structural analysis. Structural illustrations were prepared using PyMOL.

### Cell culture

HEK-293T and HeLa cells were cultured in 35 mm or 60 mm culture dishes. Cells were maintained in DMEM supplemented with 10% fetal bovine serum (FBS) and 100 U/mL penicillin-streptomycin, with fresh medium replaced every 24 h. All cells were incubated at 37 °C in a humidified atmosphere containing 5% CO_2_.

### Co-immunoprecipitation (Co-IP) assay and western blotting

HEK-293T cells were transfected with indicated plasmids using polyethylenimine (PEI) and harvested after 48 h. Cells were lysed in NP-40 lysis buffer supplemented with protease inhibitors. Clarified lysates, with 10% of the lysate reserved as input, were incubated overnight at 4 °C with anti-DYKDDDDK magnetic beads (MedChemExpress, Cat# HY-K0207). After three washes with PBST (PBS containing 0.1% Tween-20), bound proteins were eluted using 2× SDS-PAGE loading buffer, resolved by SDS-PAGE, and transferred onto PVDF membranes.

Membranes were blocked with 5% skim milk and incubated overnight at 4 °C with primary antibodies (1:1000), including rabbit anti-GFP (Cat# 50430-2-AP), rabbit anti-FLAG (Cat# 20543-1-AP), or rabbit anti-α-tubulin (Cat# 11224-1-AP) antibodies (Proteintech). After three washes with PBST, membranes were incubated with goat anti-rabbit IgG-HRP secondary antibody (1:10000; Absin, Cat# Abs20002) for 1 h at room temperature. Protein bands were visualized using BeyoECL (Beyotime) and imaged using a Tanon 5200 system.

### Antibodies and immunofluorescence imaging

Rabbit polyclonal antibodies against FLAG tag (1:1000; Proteintech, Cat# 20543-1-AP), and mouse Mocnclonal antibodies against TOM20 (1:500, ProteinTech, Cat# 66777-1-Ig) were used for immunofluorescence imaging. Secondary goat antibodies conjugated with Alexa Fluor 568 (1:1000; ThermoFisher, Cat# A11011) or Alexa Fluor 647 (1:1000; ThermoFisher, Cat# A21025) were used for fluorescence detection.

HeLa cells were fixed with 4% paraformaldehyde for 10 min at room temperature, permeabilized with 0.2% Triton X-100 in PBS for 10 min, and blocked with 2% BSA for 1 h. Cells were then incubated overnight at 4 °C with primary antibodies diluted in blocking buffer containing 2% BSA, 0.2% Triton X-100, and 0.1% Tween-20 in PBS. After three washes with PBST, cells were incubated with fluorophore-conjugated secondary antibodies for 1 h at room temperature. Nuclei were stained with DAPI, and samples were washed with PBS before mounting with antifade mounting medium. Fluorescence images were acquired using a Zeiss LSM 980 laser-scanning confocal microscope equipped with a 63× oil immersion objective lens. Image processing and quantitative analysis were performed using ImageJ software.

## Acknowledgments

We thank Drs Mingjie Zhang, Xiaotian Liu (Southern University of Science and Technology), Kaiming Zhang, Shanshan Li and Ming Li (University of Science and Technology of China) for assisting with Cryo-EM experiments and helpful disscussion. This work was supported by grants from the National Natural Science Foundation of China (32521003, 91953110, 22122703, 32170767, 32470808, 32541067), Fundamental and Interdisciplinary Disciplines Breakthrough Plan of the Ministry of Education of China (JYB2025XDXM505), the Major Frontier Research Project and the Research Funds of the Double First-Class Initiative of the University of Science and Technology of China (LS9100000002, YD9100002507, WK2490250006), the Strategic Priority Research Program of the Chinese Academy of Sciences (XDB0490000).

## Author contributions

J.N., and C.W. designed the research. J.N., M.Z., L.H., X.Z., M.L., and W.J. conducted the research. M.Z. and J.C. helped with the structural modeling. All the author analyzed the data. J.N. and C.W. wrote the manuscript. All authors approved the final version of the manuscript. C.W. supervised the research.

## Data and materials availability

All data needed to evaluate the conclusions in the paper are present in the paper and/or the Supplementary Materials.

## Supplemental Figures

**Supplementary Figure 1.**
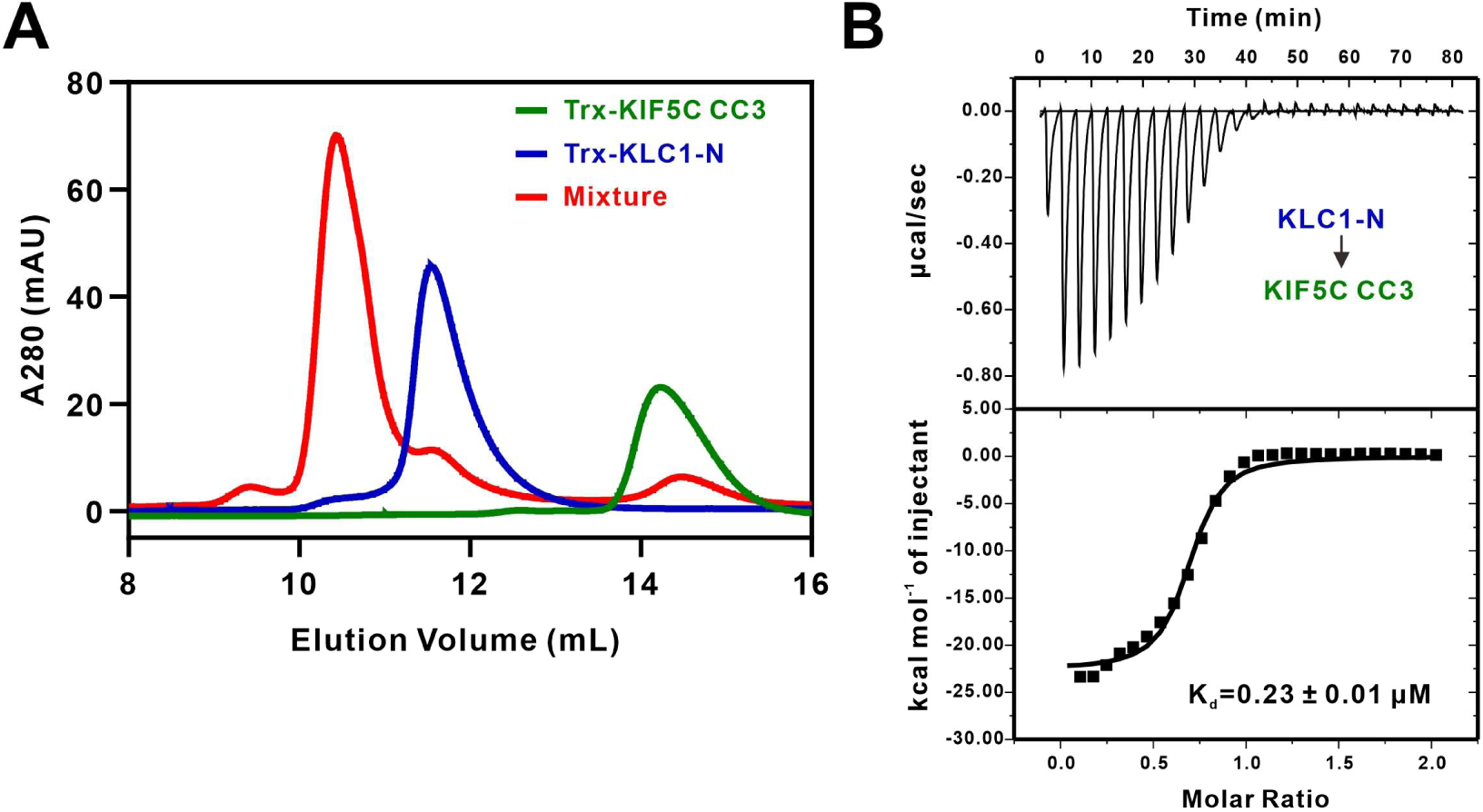
Biochemical characterization of the direct interaction between KIF5C CC3 and KLC1-N. **(A, B)** FPLC (A) and ITC (B) analyses demonstrating that the KIF5C CC3 domain directly interacts with KLC1-N with submicromolar binding affinity.

**Supplementary Figure 2.**
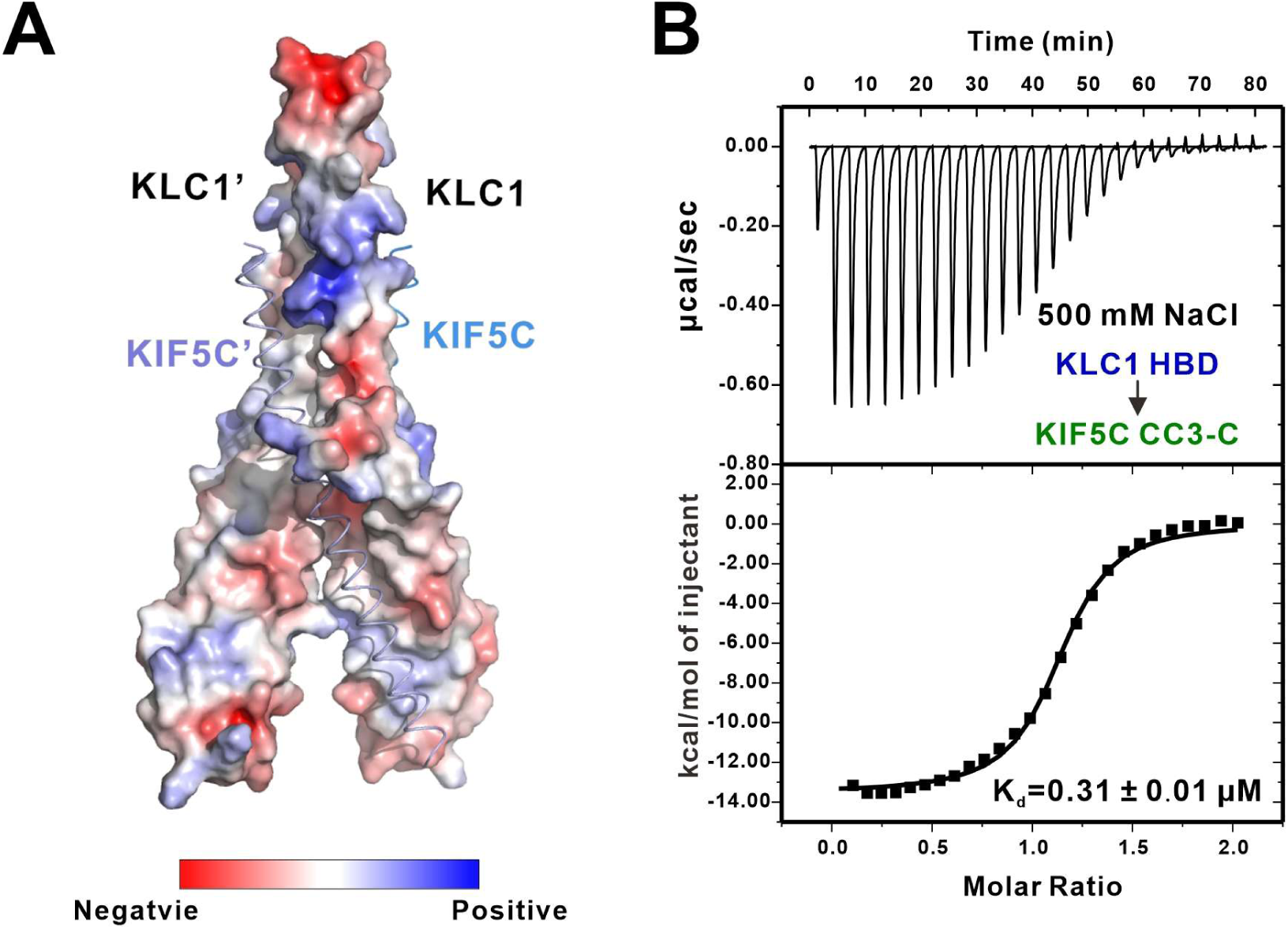
Electrostatic contributions are dispensable for KIF5C-KLC1 complex assembly. **(A)** Electrostatic surface representation of KLC1, colored from red (negative potential) to blue (positive potential), with KIF5C shown in marine blue and light blue. The KIF5C-binding interface on KLC1 lacks prominent complementary charged patches. **(B)** ITC measurement of the interaction between KIF5C CC3-C and KLC1 HBD under high-salt conditions (500 mM NaCl), showing that elevated ionic strength does not substantially affect complex formation.

**Supplementary Figure 3.**
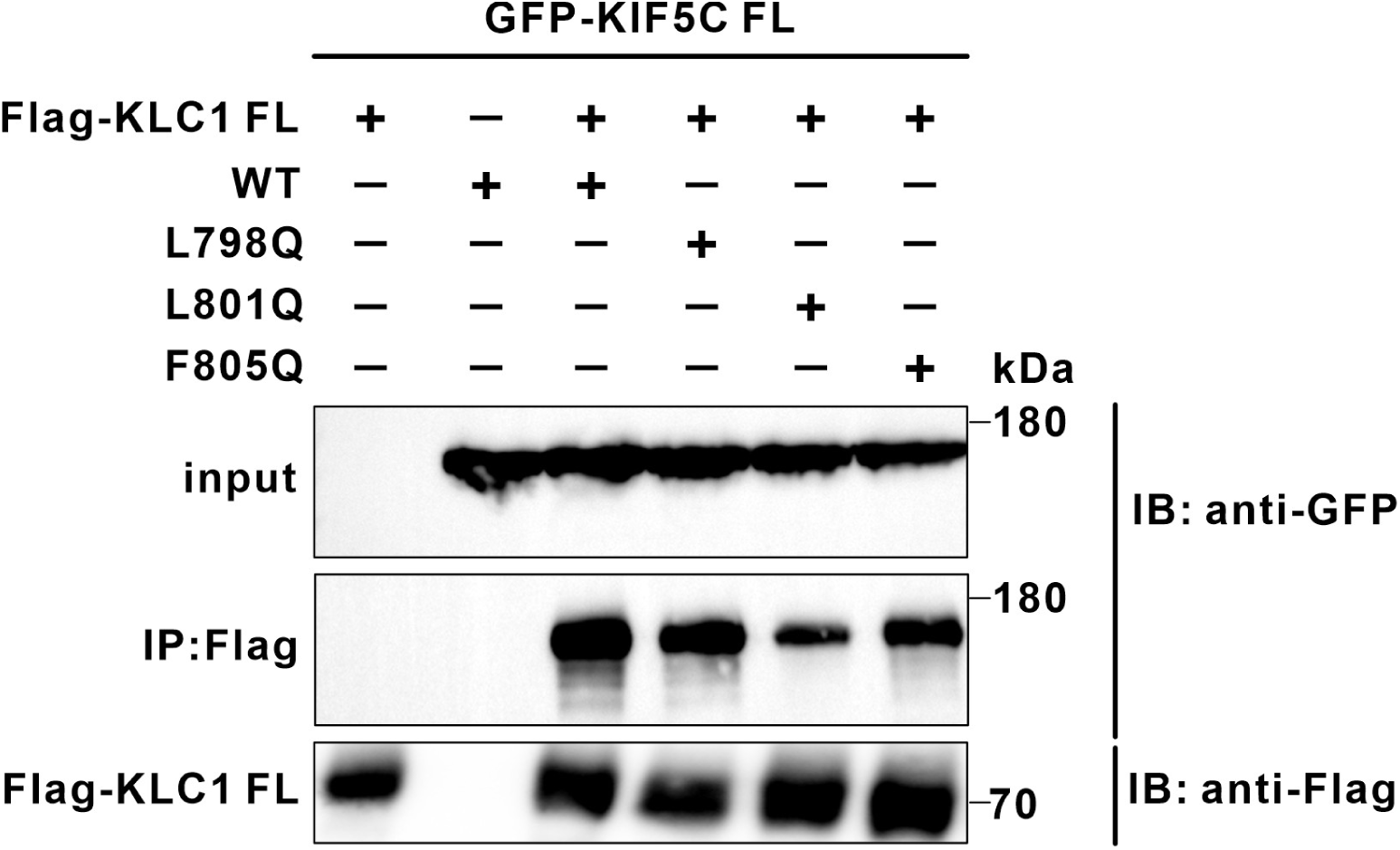
Effects of individual hydrophobic layer mutations within KIF5C on KLC1 binding. Co-IP analysis of the interaction between Flag-KLC1 and GFP-tagged KIF5C variants (wild-type or indicated single-site mutants) in HEK-293T cells. Individual substitutions within the hydrophobic interface, including L798Q, L801Q, and F805Q, partially weaken but do not abolish KIF5C-KLC1 association.

**Supplementary Figure 4.**
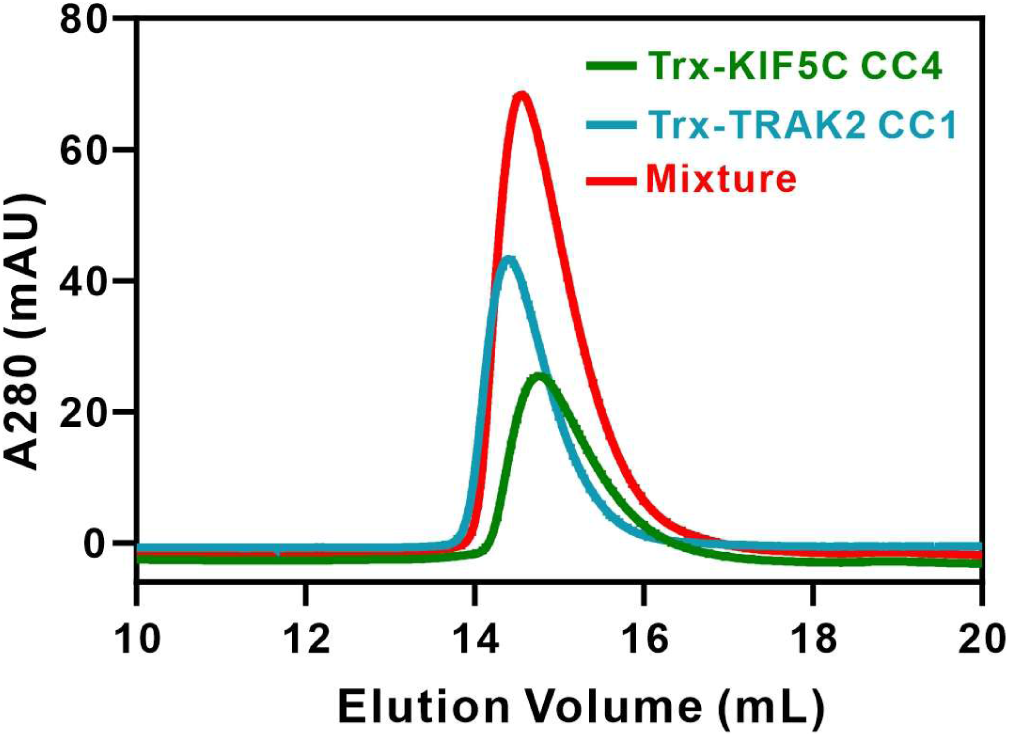
TRAK2 CC1 does not interact with KIF5C CC4. FPLC analysis showing that the TRAK2 CC1 domain does not form a detectable complex with the KIF5C CC4 domain.

**Supplementary Figure 5.**
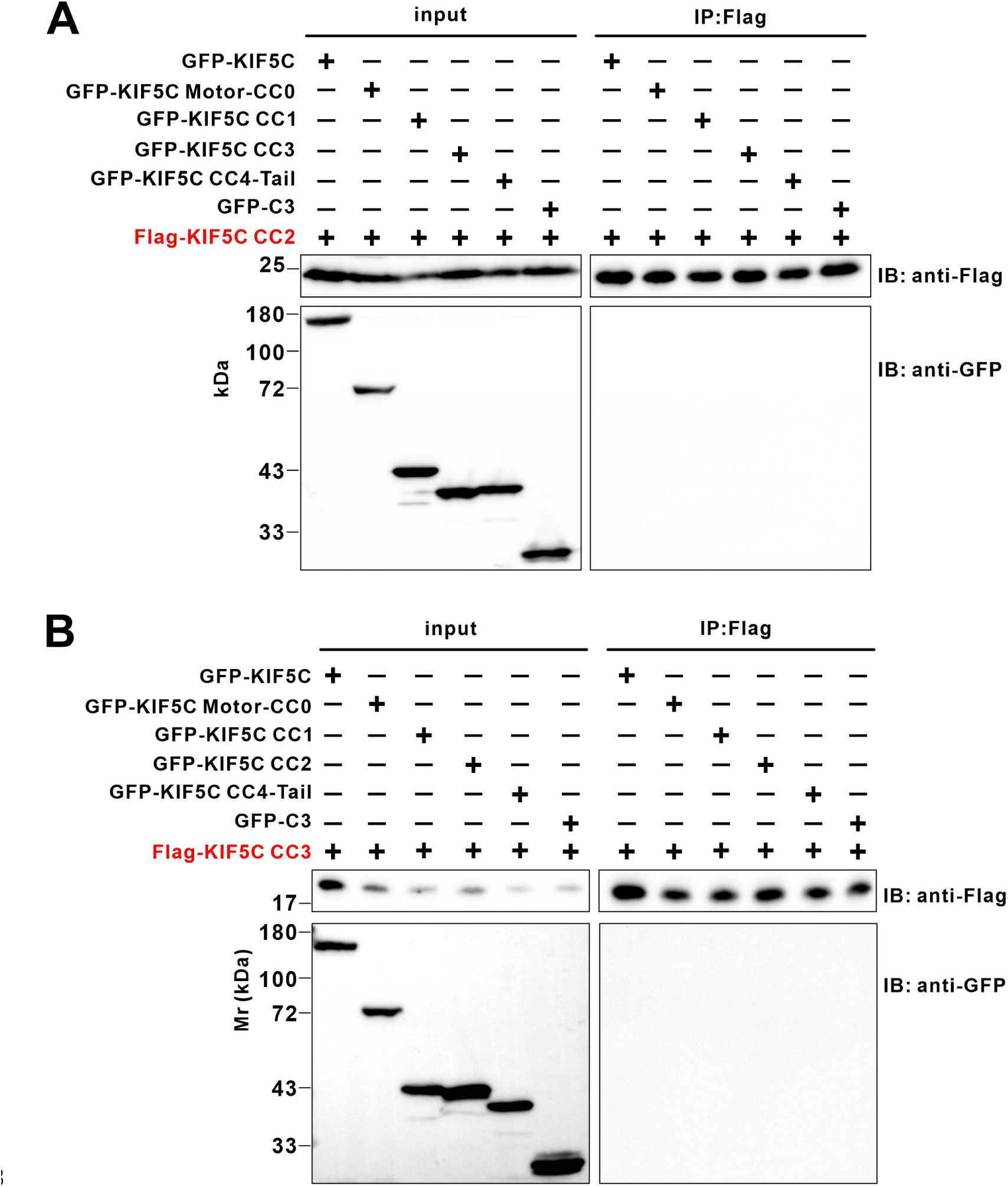
KIF5C CC2 and CC3 do not exhibit detectable intramolecular interactions with other KIF5C domains. (A, B) Co-IP analysis showing that KIF5C CC2 (A) and CC3 (B) do not detectably interact with other indicated KIF5C domains.

